# Signatures of adaptive self-organization detected in swidden societies worldwide

**DOI:** 10.1101/2025.08.04.668537

**Authors:** Sean S. Downey, Denis Tverskoi, Shane A. Scaggs, Xinyi Wu, Zaarah Syed, Jensan Lebowitz, Rongjun Qin, Stefan Thurner

## Abstract

Quantitative signatures of adaptive self-organization and the spatial scale of human management can be detected in anthropogenic landscapes. Yet, while abundant evidence for self-organization in physical and natural systems exists, there are few adaptive self-organization models that can explain coupled dynamics of human subsistence systems that have also been rigorously validated. Here, we analyze over 18,000 forest patches extracted from 18 remotely-sensed swidden mosaics from tropical and subtropical regions across the planet. We discover a power law signature of adaptive self-organization in the size of the patches and correlation distances that indicate the spatial scale of socioecological adaptation. To explain this pattern, we propose a mathematical model of swidden labor reciprocity, normative reasoning, field placement, forest disturbance and regrowth dynamics that explains the emergent spatial structure of swidden mosaics. Social self-organization can promote sustainable levels of forest disturbance when ecological dynamics such as facilitation, seed-dispersal and competition are conditioned by social dynamics, giving rise to power-law land-use patterns. We use simulations to explore the emergent properties of the model and identify two regimes: deforestation and sustainable swidden. When we analyze the relationship between the simulation results in the remote sensing data, we find that 16/18 of the study areas exhibit power-law distributions and spatial correlation patterns that match the sustainable swidden regime in the model. These findings support the hypothesis that adaptive self-organization may be a general characteristic of coupled human and natural systems.

## Introduction

Swidden agriculture (often referred to pejoratively as “slash-and-burn”) has been studied by natural and social scientists for centuries (1), but it remains a highly contested practice in scientific discourse, deepening tensions between planners and Indigenous communities—tensions that are further amplified by the escalating climate crisis. Indigenous Maya communities in Belize, like many of the 400m (2) to 1b (3) people using swidden in small rural communities around the world, rely partially or completely (2) on swidden cultivation in forests for food production – the same forests that are now seen as critical carbon sinks in the battle against climate change (4, 5). Swidden involves an annual pattern of forest clearing, burning, cultivation, and harvesting, followed by longer fallow periods when natural ecosystem processes restore above-ground biomass, species diversity, and soil fertility. Swidden farmers intervene in natural processes by creating disturbances in tropical forests that shift species composition towards those that are useful for local communities (6) and which creates opportunities for recruitment of novel species which can sustain or even enhance the overall diversity of the forest (7). Over time, swidden disturbance can create distinctive fractal-like spatial mosaics in the forest canopy that exhibit power laws distribution in patch size and correlation distances (Figure 1). This pattern is common in many natural systems (8, 9), and it has been proposed as evidence for the presence and scale of adaptive selforganization in anthropogenic landscapes such as the ancient rice terraces of Bali (10). There, synchronization of farmers’ irrigation schedules results in spatial patterns that are also visible in remote sensing imagery, power law distributions in patch sizes, and correlation distances between sets of temporally synchronized rice fields (11, 12). This process of adaptive self-organization helps guide Balinese rice agriculture to maximize rice harvests and minimize damage due to pests and water shortages (10).

**Fig. 1.**
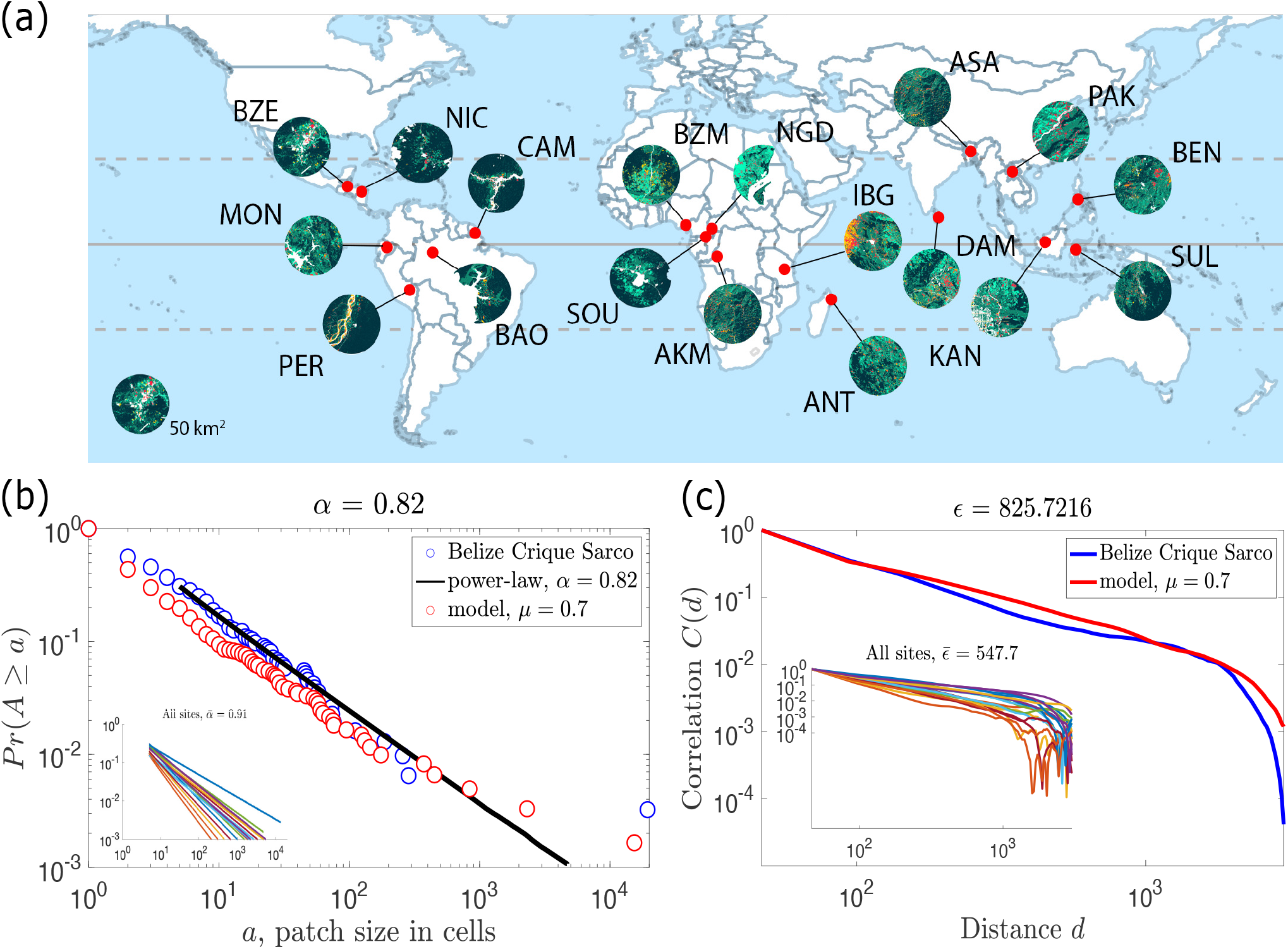
(a) Location of the 18 swidden sites that constitute our global sample and the corresponding landscape mosaics. (b) Cumulative distribution of patch sizes for Crique Sarco, Belize (blue circles) and model results (red circles). Inset: power law distribution of patch sizes across the global sample. The corresponding scaling exponents *α* are shown in the legend. (c) Correlation function *C*(*d*) versus distance *d* (in meters) for Crique Sarco, Belize (blue curve) and model results (red curve). Inset: correlation functions *C*(*d*) of raster images for all 18 swidden sites. The corresponding correlation lengths *E* are shown in the legend.

But is adaptive self-organization unique to Bali, or is it common in coupled human and natural systems (11)? Swidden agriculture provides an ideal test case for exploring this question, and the answer has important practical implications for the battle against climate change. Swidden is now most commonly observed in tropical and neotropical regions, but it also catalyzed the Neolithic expansion in Europe (13–15), was observed in the subarctic and boreal regions of Northern Europe in the 18th and 19th centuries (1), and it has gene-culture coevolutionary significance for the human species (16). However, most information about the socioecological dynamics of swidden come from ethnographic and ethnohistorical examples, where significant local ecological knowledge indicates deep cultural, historical, and linguistic connections with local forest environments. For example, our previous research in Maya communities in Belize identified a “cooperation puzzle” where more cooperative norms of swidden labor reciprocity can lead to less sustainable use of natural resources, but that the same labor exchange networks create labor dependencies between farmers which serve as a social milieu that supports sanctioning when overuse of community forests is at risk (17, 18).

We proposed that farmers negotiate their dyadic labor partnerships using a cognitive process, defined as *normative reasoning*, which helps them decide whether to provide swidden labor to a partner requesting help based on perceptions, beliefs, or understanding of the acceptability of the request (19). When we modeled this system, we identified three emergent regimes: low-intensity swidden, sustainable high-intensity swidden that maximizes ecosystem services and harvest returns, and deforestation. Theoretically, sustainable high-intensity swidden can emerge when labor reciprocity and normative reasoning are balanced: helping behavior should be significantly conditioned by normative reasoning to prevent over-harvesting, while reciprocity is necessary to prevent excessive sanctioning. Our model predicts that swidden is most productive for both forests and farmers when the balance of labor reciprocity and normative reasoning results in an intermediate scale of forest disturbance.

When we looked in remote sensing imagery for evidence of ecosystem enhancement that could result from these dynamics, we found that higher levels of biodiversity are related to intermediate levels of swidden disturbance in two Maya communities in Belize (7), and in a sample of 18 swidden societies across the planet (20). These results are consistent with Connell’s famous and controversial intermediate disturbance hypothesis (IDH) (21, 22). However, it remains an open question how social norms and swidden activities in small-scale societies that consist of tens or hundreds of households can transform forests into enhanced anthropogenic landscapes. Adaptive self-organization provides a compelling answer if low-level interactions between individual swidden farmers – including the spatial dynamics of field clearing and social dynamics related to labor reciprocity and sanctioning – can be shown to transform community forests into enhanced swidden mosaics at spatial scales that are adaptive for both human subsistence and landscape ecology. In this study, we bring together modeling and data analysis to answer this question. We analyze a unique global dataset consisting of over 18,000 forest patches extracted from 18 remotely-sensed swidden mosaics from tropical and sub-tropical regions across the planet (Figure 1, also see Material and Methods). We discover a power law signature of adaptive self-organization in the size of the patches, and correlation distances that indicates the spatial scale of socioe-cological adaptation. To explain this world-wide pattern, we extend an agent-based model of swidden labor reciprocity, normative reasoning, and sanctioning (19) by adding non-random field locations (Figure 2). In our discussion, we consider how our findings relate to intermediate-scale disturbance in swidden systems, and theoretical implications for theories of self-organization in coupled human and natural systems.

**Fig. 2.**
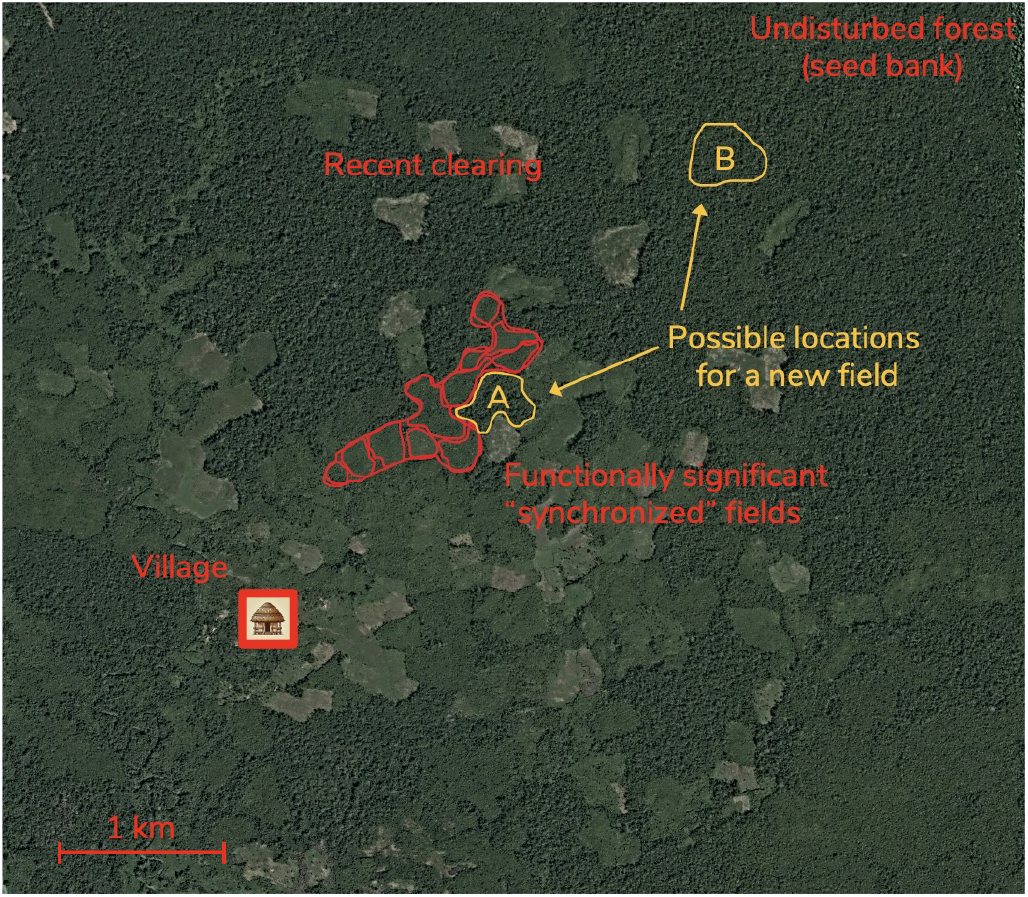
In the model, new swidden fields can be placed in a forested area adjacent to a field cleared in the previous round (e.g., Location A) if *s*_*i*_ = 1, or at a random location on the grid in undisturbed forest (e.g., Location B) if *s*_*i*_ = 0.

Our model is based on nearly 20 years of ethnographic research in Maya communities in Belize, yet the key elements of the model – social norms for labor reciprocity (23), energetic constraints (24), and forest disturbance dynamics (21, 25) – are generalizable. Swidden mosaics often develop around settlements and, after multiple cultivation cycles, adjacent forests evolve into spatially complex forest mosaics with emergent properties (26). This stems from practical farming choices such as how much forest to clear, where to plant, whom to support, and under what conditions which are typically made at the individual or household level and often influenced by social norms and traditional cultural practices. The extent and spatial characteristics of these swidden disturbances interact with ecosystem dynamics such as soil fertility recovery, seed dispersal, vegetation diversity, and forest succession to determine the emergent characteristics of the resulting swidden mosaic (20, 27).

### A spatial model of the coupled social and environmental dynamics of swidden agriculture

Our model represents a community forest similar to Figure 2, including a group of *N* swidden farming agents interacting on an *L × L* grid. Each time step *t* in the simulation approximates one year, or one annual harvest. Each grid cell in the model exists in either a forested state or a cleared state and farmers convert a set of adjacent cells to the cleared state when they cultivate it for swidden. The probability *p*_*k*_ that a cleared cell *k* will return to a forested state is governed by a facilitation model that includes two factors, seed dispersal and competition for germination (28). We assume only forests can serve as seed banks, so *p*_*k*_ depends on the fraction of surrounding cells occupied by trees within radius *R*, and other model parameters that determine the relative importance of dispersal and competition on regeneration dynamics (see the “Additional details on the model” section of the SM).

Agents in the model represent farmers who make decisions about how much forest to clear, where to place their fields, and whom to help. Their ability to achieve their desired harvests depends on social dynamics related to labor reciprocity, customary land use practices and norms, and energetic costs (19). Non-linear forest regrowth dynamics after disturbance, including ecological facilitation, affect the fractal characteristics of the emergent swidden mosaic of the simulated forest ecosystem (27). Farming agents exert a degree of control over these complex and emergent patterns through three evolving strategies. Specifically, each agent is characterized by the desired field size *x*_*i*_, the field placement strategy *s*_*i*_ *∈* {0, 1}, and an acceptable field size (sanctioning) threshold *y*_*i*_.

The field placement strategy operates as follows (see Figure 2): with *s*_*i*_ = 1, agent *i* places their new field in a forested area adjacent to a field cleared in the previous round. With *s*_*i*_ = 0, *i* searches for a new field location anywhere on the grid, subject to additional search cost, *c*_*s*_. The desired field size *x*_*i*_ determines how many forested cells the agent seeks to clear, but to do this, the agent needs help from others. However, too much labor reciprocity could also lead to over-exploitation of natural resources. How can agents overcome this? Social dynamics related to swidden labor reciprocity and social sanctioning contribute to the probability that agent *i* will help agent *j* with the request *x*_*j*_. First, helping decisions are driven by direct reciprocity: agents tend to help labor partners who have helped them in the past. Second, helping behavior is guided by normative reasoning: *i* tends to reject requests they perceive unacceptable (i.e., those in which the requested field size *x*_*j*_ exceeds their individual sanctioning threshold *y*_*i*_). Conversely, agent *i* tends to provide help in response to acceptable requests (i.e., those with *x*_*j*_ *≤ y*_*i*_). The parameter *μ∈* [0, 1] captures the relative importance of normative reasoning versus reciprocity in helping decisions. Helping behavior also has an associated cost of helping. In this way, the average sanctioning threshold 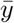 can be interpreted as a sanctioning norm.

The individual harvest *x*_*a,i*_ is defined as the number of cells cleared by agent *i* and is determined by the clearing request *x*_*i*_, the number of helpers, and the number of available forested cells. The individual payoff *π*_*i*_ is defined as the harvest obtained minus helping and search costs. Each round, agents have the opportunity to revise their strategies. The desired field size *x*_*i*_ is updated via myopic optimization, in which the agent chooses the value of *x*_*i*_ that maximizes their payoff, assuming that others keep their strategies the same as in the previous round (29–31). Myopic optimization reflects the bounded rationality of agents who are able to maximize their payoff but are unable to predict changes in the behavior of others. The acceptability threshold *y*_*i*_ and the field placement strategy *s*_*i*_ are updated using payoff-biased imitation with innovations. That is, agents are more likely to adopt strategies from others who achieved higher payoffs, with occasional experimentation to explore new options. In real swidden communities, decisions about where to locate a field, how much to clear, which other farmers to help and which to reject, are quite complicated involving discussions with family, friends and other farmers, as well as social capital and customary agroecological knowledge. We do not capture this level of detail and instead model this decision in a very simple way to investigate whether the system can adapt to the co-evolving characteristics of the forest. In one important way the model is similar to real swidden communities: farming agents do not have comprehensive information about spatial characteristics of the grid, or about the underlying ecological parameters governing forest regrowth. Overall, each time step proceeds as follows:

1. **Forest ecology**. Each empty cell becomes forested with probability *p*_*k*_. The amount of new biomass 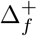is equal to the number of cells that regenerate.
2. **Field planning**. Each agent *i* determines the desired field size *x*_*i*_ via myopic optimization and selects a location based on their field placement strategy *s*_*i*_.
3. **Helping behavior**. For all agents *i* and *j*, calculate the probability *q*_*i,j*_ that *i* (having their acceptability threshold *y*_*i*_ and the current history of labor exchange interactions) will help *j* with the request *x*_*j*_.
4. **Harvesting**. Each agent *i* receives a harvest *x*_*a,i*_ based on their request *x*_*i*_, the number of agents that provided help *i*, and the number of available forest cells.
5. **Payoff calculation**. For each agent *i*, calculate their payoff *π*_*i*_ accounting for individual harvest, helping costs, and search cost.
6. **Grid update**. Clear the corresponding number of forest cells for each agent at the chosen location. The remaining fraction of forest *f* is recorded.
7. **Helping history updates**. The history of labor exchange interactions is updated for all agents.
8. **Strategy update**. Each agent *i* updates their sanctioning threshold *y*_*i*_ and field placement strategy *s*_*i*_ using payoff-based imitation with innovations.

Each model simulation is initialized with parameters *L* (grid size), *c*_*s*_ (search cost), *N* (number of agents), *R* (seed dispersal radius), and *μ* (relative importance of normative reasoning). Agents are assigned randomized initial field placement strategies *s*_*i*_, field size strategies *x*_*i*_, and sanctioning thresholds *y*_*i*_. For a formal mathematical model description and the default parameter values, see the “Additional details on the model” section of the SM.

## Results

### Quantitative signatures of self-organization

In our model, interactions among labor reciprocity, normative reasoning, field size, field placement, and forest regrowth dynamics affect harvest returns, the emergent spatial structure of swidden mosaics, and the emergence of power-law distributions and spatial correlation patterns. Simulations in Figure 3 reveal two distinct regimes, deforestation and sustainable swidden. In the deforestation regime, strong labor reciprocity and lesser importance of normative reasoning enable agents to use their labor exchange network to clear larger fields, leading to excessive forest exploitation and environmental degradation (Figure 3a). In contrast, the sustainable swidden regime prevents deforestation and leads to the emergence of forest mosaics with scale-free properties and adaptive labor group dynamics (Figure 3b, d-e). The sustainable swidden regime is observed when the relative importance of normative reasoning *μ* exceeds a threshold, that is determined by demographic and environmental parameters. Specifically, the range of *μ* that supports the sustainable swidden regime expands with longer-range seed dispersal *R* (Figure 4a), and lower population density (Figure 4b). In the sustainable swidden regime, labor reciprocity and normative reasoning respond dynamically to social and environmental conditions. Two indicators of adaptive self-organization and the spatial scale of coupled human and natural systems are power law distributions of patch sizes and correlation lengths (11). We assess whether swidden exhibits these characteristics by first defining a synchronized patch as the contiguous area of the same land class irrespective of farmer. We then analyze whether these distributions resemble power laws in both the simulation model and in the remote sensing data. In the simulation, the deforestation regime does not exhibit power laws because it consists of a single cleared patch. However, we cannot reject a power-law model in more than 90% of the simulation runs sampled from the sustainable regime using the Kolmogorov-Smirnov goodness-of-fit test (see Table S4 in the SM). A similar scaling pattern is found in the remote sensing data from 18 swidden sites across the globe: in 16 out of 18 cases, power-law behavior cannot be ruled out (see Figure 1 and Table 1). This convergence between model and data is supported by their comparable average scaling exponents: 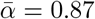 in the model and 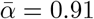 in the remote sensing data.^1^ Next, we examine correlation lengths *E* in the remote sensing data (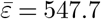, *σ*_*ε*_ = 252.3, see Table 1 for details) and in the simulations (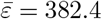, *σ*_*ε*_ = 194.3). In both cases, spatial correlations exhibit a pattern of decay with distance that also resemble a power law (see Fig. 1c, inset and Figure 3b,d-e). The shorter correlation lengths in the model compared to the data are due to spatial factors not represented in the model — such as clustering fields near villages or rivers, avoiding areas due to terrain or soil quality, or respecting land demarcations. Overall, the correspondence between indicators of adaptive self-organization in our swidden model and remote sensing data stimulate further investigation of the underlying mechanisms and dynamics that support the sustainable swidden regime.

**Fig. 3.**
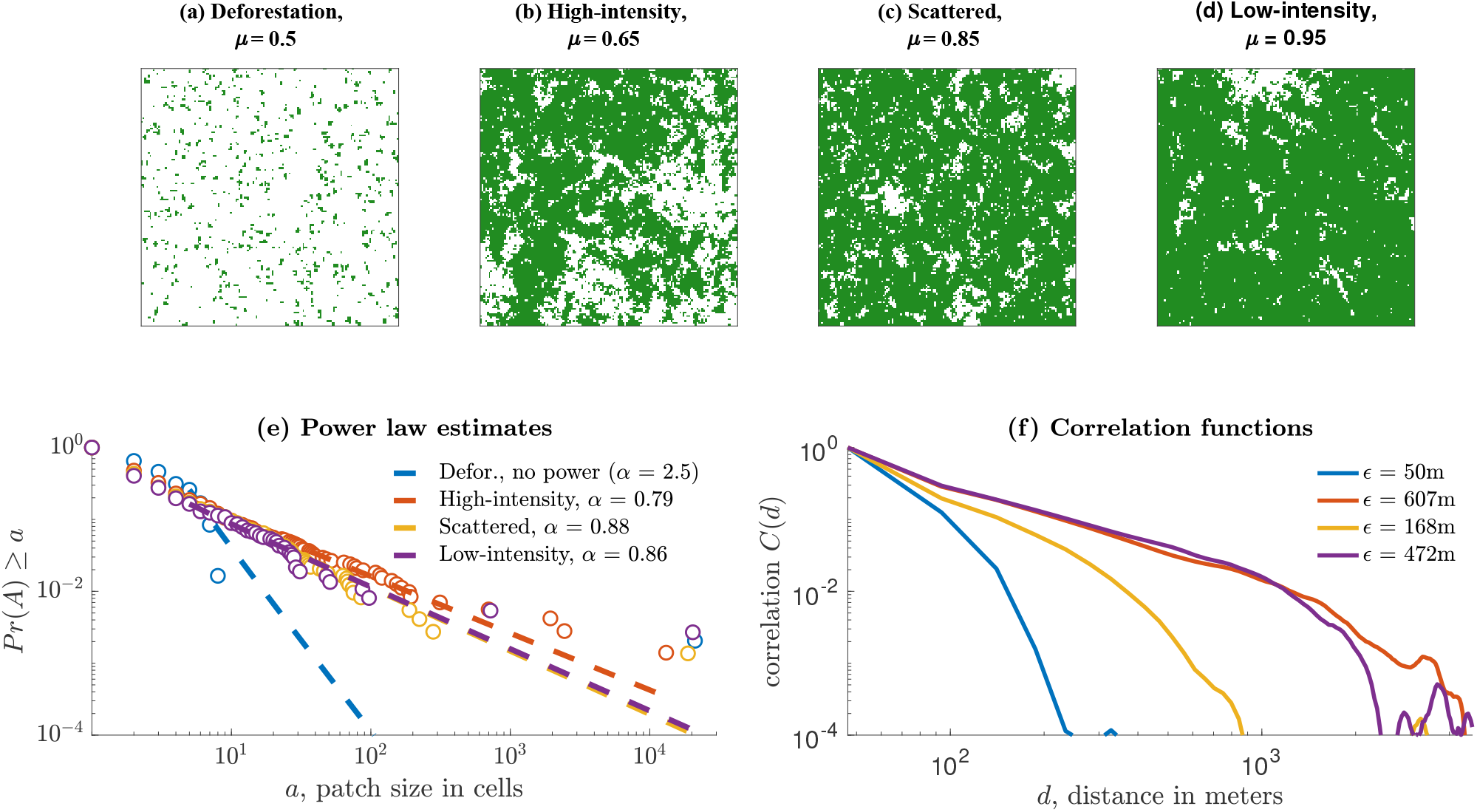
Panel (a) illustrates the spatial and temporal pattern that characterizes the deforestation regime, and panels (b-d) illustrate three characteristic patterns that occur in the sustainable swidden regime. Each panel displays the forest landscape either at the end of the simulation run or in the case of deforestation, 10 rounds before complete deforestation. Panel (e) shows the distribution of patch sizes *a*, and panel (f) shows spatial correlations *C* as functions of distance (in meters). For additional information about these simulations see section “Additional results from the main model” of the SM.

**Fig. 4.**
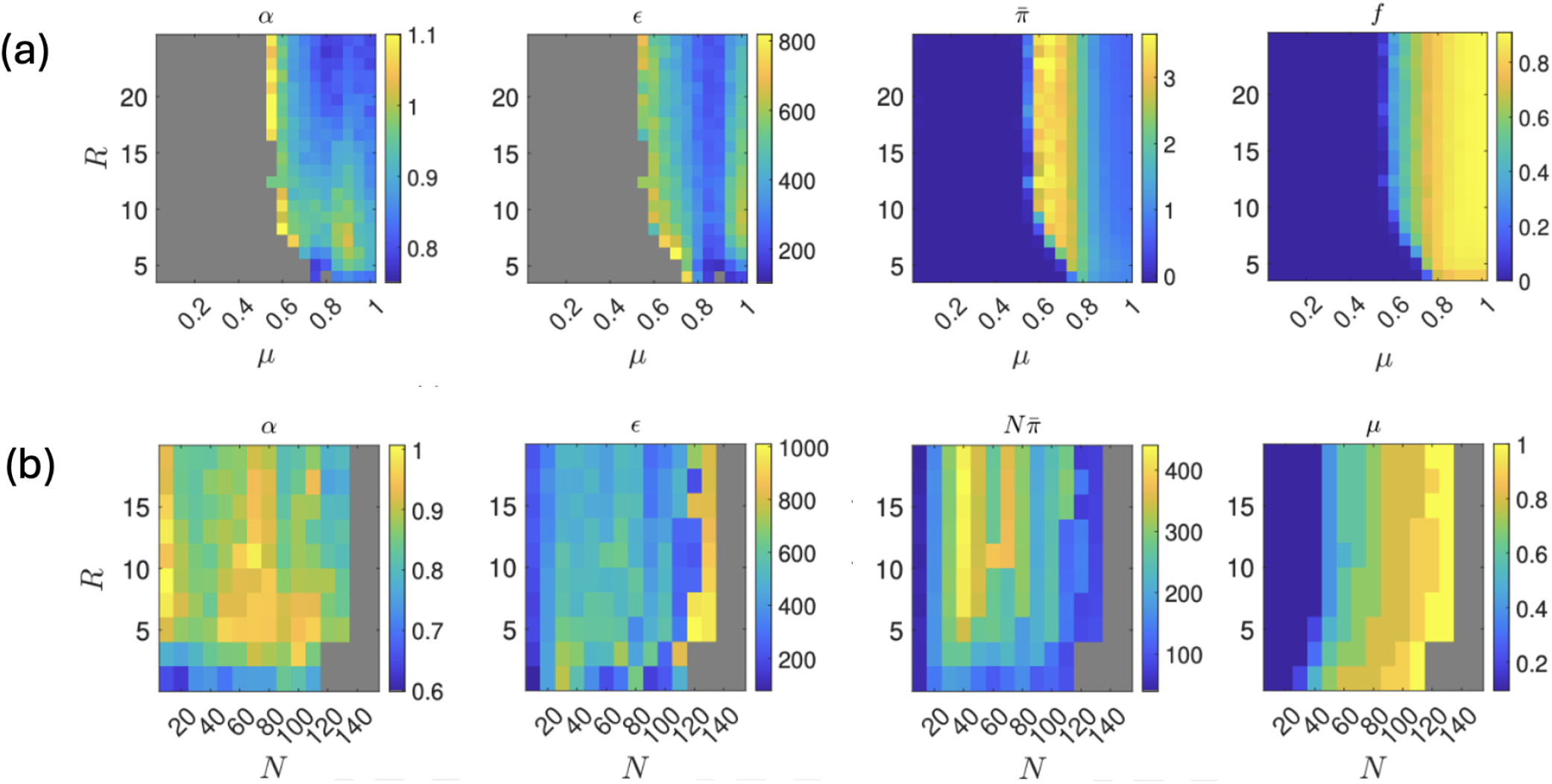
(a) Effects of the social parameter *μ* (representing the relative importance of normative reasoning) and the ecological parameter *R* (the seed dispersal radius) on key model outcomes: the remaining fraction of forest *f*, the individual payoff *π*, the scaling exponent *α*, and the correlation length *ε*. For each (*μ, R*) combination, 100 independent simulation runs were performed. Each run consists of 1000 rounds, with outcome measures averaged over the final 250 rounds. All other parameters are set to their default values. (b) The effects of group size *N* and the ecological parameter *R* on group payoff 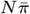, the scaling exponent *γ*, and the correlation length *ξ*. Varying the parameter *μ* represents an additional layer of adaptation for the swidden group. To illustrate this, for each (*N, R*) combination, we select the value of *μ* that enables the group to achieve a high harvest while maintaining reasonable levels of deforestation risk. Therefore, for each (*N, R*) combination, 60 independent simulation runs were conducted for each value of *μ ∈ {*0.1, 0.2,…, 1.0*}*. Among these, the value of *μ* that yielded the highest average individual payoff—while also resulting in a sustainable swidden regime in more than 60% of runs—was selected. The reported values represent averages over the corresponding 60 runs for each *N, R* pair. Each run consists of 1000 rounds, with outcome measures averaged over the final 250 rounds. All other parameters are set to their default values.

**Table 1.** Summary of swidden sites. The table reports the scaling exponents *α* of the patch size *a* distributions, with standard errors *θ* given in parentheses. Also shown are p-values from the Kolmogorov–Smirnov goodness-of-fit test, where *p >* 0.1 indicates that the power-law model cannot be rejected, and the estimated correlation length *ε*. Patches are defined as contiguous areas of the same land class in the remote sensing image grid, using a Von Neumann neighborhood for contiguity. Patch sizes are measured in units of the area of a single grid cell. The discrete power-law model is fitted with a minimum patch size *a*_*min*_ corresponding to five grid cells (approximately 1.1 hectares). Standard errors *θ* are calculated using a bootstrap procedure.

| Site | Country | $n_{a \geq a_{min}}$ | Scaling exp. $\alpha$ ($\theta$) | p-value | Corr. length, $\varepsilon$ (m) |
| --- | --- | --- | --- | --- | --- |
| Crique Sarco | Belize | 95 | 0.82 (0.10) | 0.42 | 826 |
| Borneo | Indonesia | 131 | 0.79 (0.09) | 0.44 | 400 |
| Boa Vista | Brazil | 36 | 0.95 (0.30) | 0.34 | 394 |
| Banzoum | CAR | 121 | 0.91 (0.11) | 0.70 | 972 |
| Nganda | DRC | 91 | 0.73 (0.10) | 0.75 | 637 |
| Souanke | DRC | 76 | 0.81 (0.12) | 0.90 | 492 |
| Mondayacu | Ecuador | 134 | 0.83 (0.08) | 0.17 | 218 |
| Camopi | French Guiana | 57 | 1.03 (0.28) | 0.57 | 858 |
| Asananggre | India | 100 | 1.32 (0.24) | 0.71 | 168 |
| Sulawesi | Indonesia | 88 | 1.14 (0.18) | 0.44 | 339 |
| Pakkeng | Laos | 167 | 0.80 (0.07) | 0.03 | 717 |
| Antalaha | Madagascar | 164 | 0.97 (0.10) | 0.84 | 522 |
| Arang Dak | Nicaragua | 34 | 0.82 (0.20) | 0.66 | 557 |
| Akamkpa | Nigeria | 114 | 1.06 (0.14) | 0.78 | 248 |
| Nuevo Eden | Peru | 54 | 0.58 (0.09) | 0.65 | 779 |
| Benli | Philippines | 121 | 1.22 (0.18) | 0.56 | 212 |
| Dambana | Sri-Lanka | 164 | 0.78 (0.08) | 0.09 | 756 |
| Ibingu | Tanzania | 140 | 0.88 (0.10) | 0.35 | 764 |

### Mechanics of self-organization

To investigate the coupled social and ecological dynamics of our model, we simulate how the structure of the swidden mosaic changes in response to changes in the relative importance of normative reasoning *μ* with a fixed seed dispersal radius (*R* = 13, Figure 3). At the lowest values (*μ ≤* 0.55, Figures 3a and S2c in the SM), labor reciprocity quickly drives the forest past its carrying capacity resulting in a single large deforested patch. However, with a moderate degree of normative reasoning, swidden cultivation can occur sustainably and harvest levels are maximized. Between (0.55 *μ* 0.75), labor reciprocity is tempered by normative reasoning, farming intensity decreases, and the forest exhibits bifurcation dynamics that are sensitive to abrupt transitions from sustainable outcomes to deforestation (see Figure S2c in the SM). At the highest levels, (*μ >* 0.75), swidden activity occurs sustainably, but harvests levels are low. In the sustainable swidden regime, there are three discernible spatial patterns that emerge at different levels of *μ*. To illustrate these differences, we simulate at three different levels of *μ* which we label high intensity swidden, scattered swidden, and low intensity swidden.

High intensity swidden (*μ* = 0.65, Figure 3b) is characterized by high levels of spatial synchronization of the swidden mosaic, reflected in a large correlation length *ε*, that is driven by moderately sized fields, synchronized swidden patches, high harvest returns and payoffs, and high biomass productivity. This outcome arises because intermediate values of *μ* promote moderate sanctioning norms and medium-sized fields. Field synchronization means large contiguous swidden patches could form, but patch size is constrained by dynamic field placement strategies *s*, which balance field synchronization and pioneer farming. This adaptive behavior enhances individual payoffs, as the benefits of searching for new harvest areas outweigh the associated costs, and aggregating moderately sized fields into larger swidden patches can maximize biomass recovery rates due to the scale-dependent regrowth dynamics of the simulation. The distribution of swidden patches form a power law (Figure 3e) due to the mix of medium and large swidden clearing and medium and large forest patches. The shape of the correlation function (Figure 3f) reflects the integration and maintenance of long-range landscape connectivity required for seed dispersal from forest seed banks to swidden clearings and the energetic benefits of medium-range synchronization of swidden fields.

Scattered swidden (*μ* = .85, Figure 3c) is defined by an anthropogenic state that balances connectivity of forested patches and synchronized swidden patches. It is characterized by smaller labor groups and smaller fields because more stringent sanctioning occurs with a higher importance or normative reasoning and small swidden patches that appear randomly scattered throughout the forest. Large synchronized swidden patches do not occur because fields are relatively small and agents establish new fields away from their current clearings with moderate frequency. The largest patch is a single forested component, which is interspersed with smaller synchronized swidden patches. Scattered swidden exhibits adaptive behavior because the benefits of pioneer locations are balanced against the lower search costs of synchronizing with existing fields, so the distribution of synchronized patches forms a power-law distribution and the correlation lengths are lower than in high intensity swidden (Figure 3e,f).

Low intensity swidden (*μ* = .95, Figure 3d) occurs as normative reasoning curtails or eliminates most group farming activity, which significantly limits field sizes. As a result, farmers have little incentive to place new fields away from their recent ones, as doing so would incur unnecessary search costs, leading to the formation of small individually cleared synchronized swidden patches. The distribution of patches also forms a power law distribution. In this example (Figure 3d), the two largest patches are a forest component and a swidden patch which could be considered outliers to the power law distribution; however, swidden synchronization is driven by the co-occurrence of large forested areas and small cleared patches that form stable swidden areas for each agent, that result in a high mean correlation distance. Overall, low intensity swidden occurs sustainably, but the stringent sanctioning norm limits the larger clearings so the size of synchronized swidden patches is smaller, resulting in less overall biomass productivity. The low intensity swidden regime relies on individual labor rather than group labor, so it does not produce large harvests, and it is unlikely to be economically or calorically viable for long periods of time.

In summary, Figure 3 shows how interactions among field size, sanctioning norms, and field placement strategies can result in sustainable or unsustainable outcomes. At least three characteristic spatial patterns can appear in the sustainable swidden mosaics, all of which result in comparable values of *α*, but different *E* (see Table S4 in the SM). Additional information on how social and ecological factors interact dynamically to produce observed swidden mosaics can be found in Table S3 and Figure S2 in the SM.

### Adaptability of self-organization

We investigated the adaptability of the coupled dynamics of labor reciprocity, normative reasoning, and clearing strategies to changes in ecological and demographic conditions by running simulations with different population sizes, forest regeneration capacities, and stochastic shocks. We found that the sustainable swidden regime emerges under a wide range of conditions. For example, Figure 4a illustrates how the social parameter *μ* can modulate the dynamics underlying swidden labor exchange behavior to maintain optimal individual-level payoffs 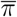 and a relatively high fraction of remaining forest *f* when decreasing seed dispersal radius *R* reduces the probability of seed recruitment in a cleared patch. Similarly, Figure 4b illustrates how *μ* can maintain group level payoffs 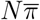 by adapting to changes in both seed dispersal *R* and population levels *N*. In both examples, the simulations explore adaptation to smooth, continuous changes in the fitness landscape. We also explored how stochastic exogenous shocks, simulating the effects of landscape-scale disturbances, for example how a forest fire might affect swidden dynamics (see Figure S3 in the SM).

In most cases, the effects on payoff and forest recovery of punctuated, large-scale disturbance simulations are similar to those of increasing population and inhibiting seed dispersal dynamics and forest regeneration. This occurs because individual (agent-level) preferences related to helping behavior *μ* ultimately lead to the evolution of emergent sanctioning norms that can adapt to a dynamically changing ecology. Interestingly, however, the values of the scaling exponent *α* remain remarkably consistent across varying ecological and demographic conditions, suggesting that swidden groups are able to maintain levels of forest disturbance that give rise to a scale-free distribution of spatial land-use patches. In contrast, the correlation length *E* is more sensitive to these changes. For example, it tends to be lower at low population densities, because a finite swidden labor pool constrains the ability to clear large patches, which in turn limits spatial synchronization over larger distances (see Figure 4b). Conversely, large-scale environmental disturbances, such as hurricanes or wildfires, can increase correlation lengths (Figure S3 in the SM).

Overall, swidden agriculture is resilient to a wide range of environmental conditions and shocks, helping to mitigate negative consequences by adjusting land use patterns and swidden labor exchange network dynamics. Adaptation occurs as individual decisions about where to establish new fields, how much land to clear, and whom to help in field clearing and cultivation interact with ecosystem dynamics leading to emergent landscape-scale properties that optimize harvests.

## Discussion

### Adaptive self-organization in swidden systems

We propose that the adaptive self-organizing model of swidden agriculture presented and validated against a global dataset of 18 swidden societies is an important advance in theoretical understanding of swidden systems. Previous theoretical approaches based on models of demographic growth, limits imposed by carrying capacity, and reduced fallow length (e.g., (32, 33)) fail to capture the social and ecological dynamics that are most important at the village scale, where adaptive behavior is more likely to occur. For example, many models of swidden are built explicit assumptions of carrying capacity (i.e., Malthusian dynamics), and *K* is defined as the total number of individuals that can be supported on a landscape at a given growth rate *r*. Others models prioritize the importance of technological enhancements *τ* which increase carrying capacity (i.e., *τ K*)(34). Such models, however, typically ignore localand mesoscale dynamics and locally enhanced or depressed levels are averaged into a single parameter *K*. Another important approach for analyzing swidden systems – common in field-based studies like anthropology and geography – is to use ethnographic, ethno-scientific, or behavioral ecology methods to document Indigenous and customary knowledge systems (35), for example by classifying economically useful species (6), documenting customary ecological knowledge and measuring harvest patterns and returns (36), by examining how social norms can help to manage common property resources (37), or by identifying alignments between ritual practices and local ecological dynamics (38). In these studies, swidden societies are often seen as having intimate, co-evolved, and/or symbiotic relationships with their local environments where social norms and practices mimic natural ecosystem processes (39), or where agroforestry practices have been developed to support and enhance natural processes (40). Such accounts have identified traditional ecological knowledge that suggests conservative environmental ethos. But the geographic and cultural specificity of these studies makes them difficult to generalize or apply in policy contexts. Our model is informed by longterm ethnographic research in Maya communities in Belize, as well as this broader literature on the traditional ecological knowledge of swidden agriculture. Our results complement the sustainable conclusions of many of the ethnographic studies by explaining how local-scale farming activities that are informed by TEK can enhance large-scale environmental processes. Specifically, our model leverages theoretical insights from complex systems theory to provide a mechanistic explanation for how TEK applied by small communities of farmers at the patch level can generate common emergent patterns at the forest/landscape scale that can maintain large-scale ecosystems at a bi-stable high intensity swidden state. But how exactly does this occur? A third approach (IDH) provides a missing link between forest disturbance and ecosystem enhancement.

The intermediate disturbance hypothesis (IDH) has been proposed as a useful heuristic model for understanding the sustainability of swidden agriculture (41–43). The hypothesis suggests that ecosystems require an intermediate level of disturbance to exhibit the highest level of productivity and diversity: too much disturbance will overwhelm the system’s ability to recover, while too little disturbance will reduce species diversity and productivity (21). The relationship to swidden is intuitive because patch clearing for swidden fields disturbs forest canopies in a similar manner that natural disturbances such as tree-falls of forest fires. Recently, statistical evidence supporting this claim has begun to accumulate (7, 20). In contrast to carrying capacity or technological enhancement models, the IDH models carrying capacity as a function of the spatial and temporal scales of disturbance. Swidden cultivation is a socioecological system that creates localized disturbances capable of enhancing the productivity of the landscape by creating ecological niches that favor plant recruitment and diversity, so under certain conditions, it can stimulate marginal increases in the overall carrying capacity of the landscape and create a win-win solution for farmers by increasing harvest levels, and for the forest by increasing biodiversity and productivity (see Figure 5).

**Fig. 5.**
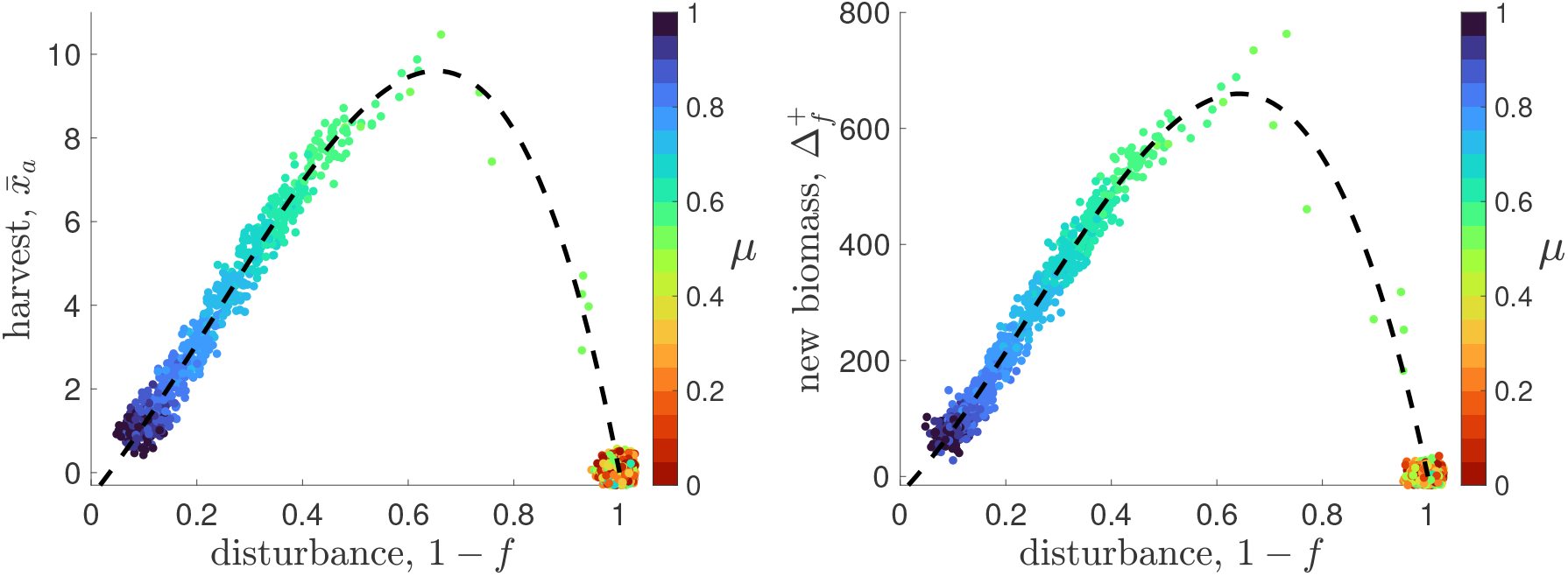
Simulated long-term disturbance curves showing the dependence of the equilibrium average harvest 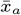 (left) and the new biomass 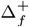 (right) on the disturbance level (1 *− f*). Each point represents the outcome of a single simulation run, with values averaged over the final 250 of 1000 total rounds. Point colors indicate the relative importance of normative reasoning *μ*. Results are shown for 100 runs at each value of *μ ∈ {*0, 0.05,.., 1*}*. All other parameters are set to default values.

However, forests are complex systems (44) consisting of interacting geology, soil, plant, animal, and climatic elements subject to non-linear interactions across temporal and spatial scales, including in forest succession dynamics (27). Such systems are susceptible to regime shifts and collapse, which can be seen in our model to the right of the bifurcation point *f* = 0.6 (see Figure 5). The adaptive self-organizing swidden model we present here provides a theoretical explanation for how social norms related to reciprocity and normative reasoning can help small-scale societies – who have limited environmental, social information and capacity for labor – to maintain forests to the left of the bifurcation point. In the model, individual (or household-level) decisions about field size, field placement, and who to help create self-organizing dynamics that support an emergent sustainable swidden regime by synchronizing swidden fields to create spatially correlated landscape mosaics at a spatial scale that optimizes underlying ecosystem disturbance and regrowth dynamics. When this occurs successfully, it corresponds with high intensity swidden in our model where enhanced forest productivity and higher harvest returns occurs. However, the model is non-linear and the high-intensity swidden regime is a bi-stable equilibrium point, so continuous adaption is necessary and alternative states can occur unpredictably. When the entire set of sustainable and deforestation states are considered, we find that swidden disturbance at an intermediate scale – corresponding to high-intensity swidden – enhances ecosystem productivity in line with the IDH, ultimately resulting in the highest harvest returns and payoffs, supporting results reported elsewhere (7, 20). Taken together, our adaptive self-organization model of swidden agriculture explains how social dynamics mediate these complex socioecological dynamics and, in doing so, it provides an important alternative to the demographic and ethnographic frameworks for understanding swidden agriculture.

### Universality of patch correlation in coupled human and natural systems

The Balinese subaks provide a prototypical example of adaptive self organization that optimizes irrigated rice harvests by matching the scale of spatial synchronization of irrigation schedules with pest and water induced stresses. However, the continuous 1,000+ year history of the subak system raises questions about whether the quantitative signatures of self-organization detected in remote sensing imagery are unique to Bali. So, is adaptive self-organization common in coupled human and natural systems worldwide as the authors of (11) hypothesize? Neither the rice terraces or swidden mosaics exhibit evidence of topdown natural resource management institutions, instead both are examples of commons management (10, 45). However, unlike the highly engineered and postcard-worthy (11) irrigated rice terraces and unique culture of Bali, swidden entails minimal physical infrastructure, appears to be more globally widespread, and has likely been an important subsistence practice for much of the Holocene.

Our analysis of swidden mosaics from 18 small-scale and Indigenous societies samples are sociocultural diverse and they represent a wide range of geographic and environmental variability. Similar to the Bali study, we find quantitative signatures of adaptive self-organization in the size distribution and spatial correlation of synchronized swidden patches. It is notable that a comparison of the empirical scaling exponents and correlation distances both appear to generate similar quantitative signatures of adaptive self-organization (Figure 6). Slightly lower scaling exponents and increased variance in the correlation distances in swidden forests reflect more tightly constrained socioecological dynamics in Balinese rice terraces (irrigation and rice pests) than in global swidden forests (labor dynamics and seed recruitment).

**Fig. 6.**
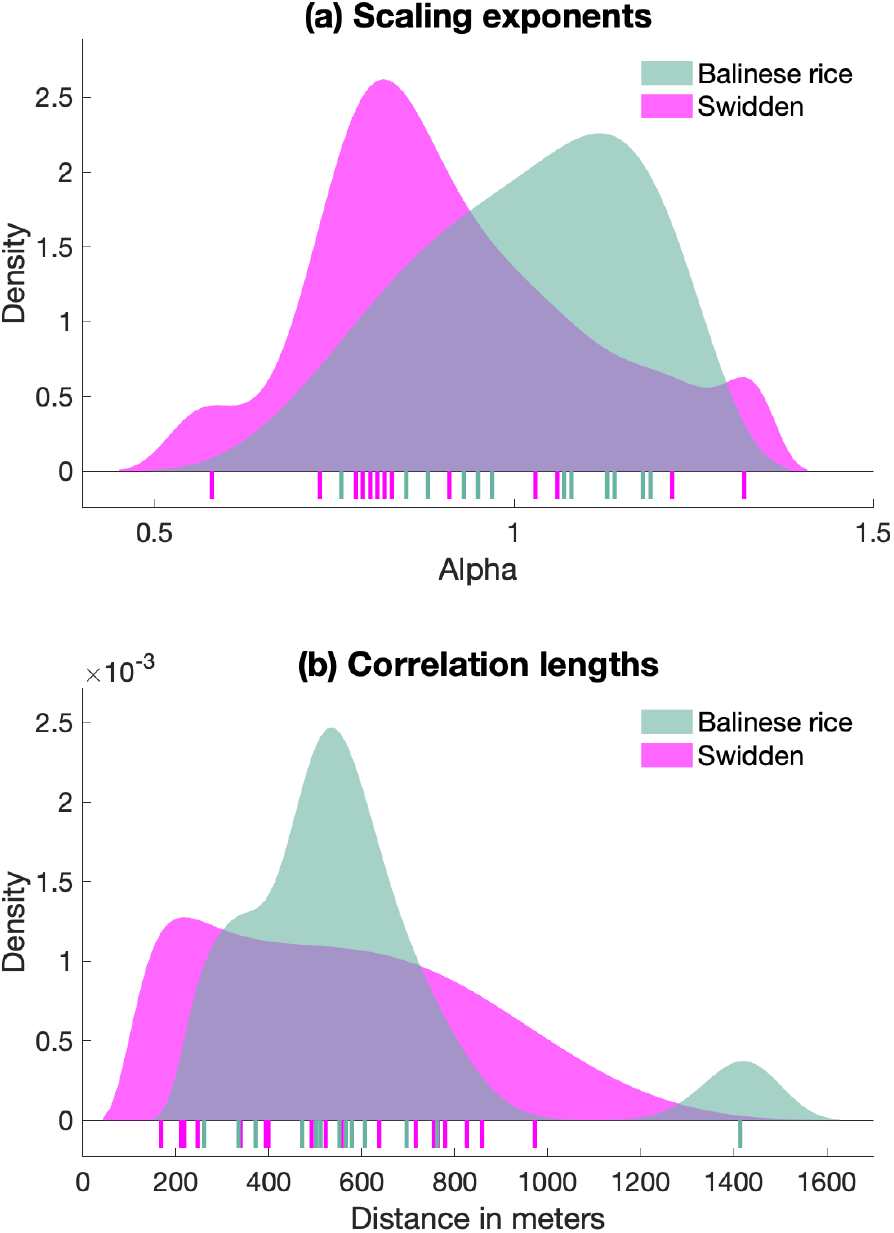
Comparison of (a) the distributions of estimated scaling exponent *α* and (b) correlation length *ε* in swidden and Balinese rice systems. The mean scaling exponents for swidden (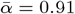, *σ*_*α*_ = 0.18) is slightly lower than for Balinese rice (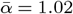, *σ*_*α*_ = 0.14, *t* = *−*1.84, *df* = 28.9, *p* = 0.08), indicating that the there are more large patches in swidden mosaics than rice terraces. The correlation length in swidden (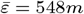, *σ*_*ε*_ = 252*m*) appeared to be comparable with that in Balinese rice (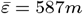, *σ*_*ε*_ = 284*m, t* = *−*0.40, *df* = 24.1, *p* = 0.70). Estimates of *α* and *ε* for Bali are transcribed from supplementary information in (11).

The model of swidden cultivation that we developed includes fundamental coupled social and ecological dynamics related to labor reciprocity, commons management, disturbance from swidden cultivation, and forest regrowth that should be generalizable to many known instances of swidden agriculture. In the model, the harvest levels serve as an indirect signal of environmental state, enabling adaptive decision-making in the absence of direct observation of macro-scale ecosystem productivity. Farming agents adapt to declines in harvest returns by limiting labor group sizes and areas harvested to allow forest recovery and sustainable cultivation. They increase swidden activity when environmental and social conditions permit by enabling larger labor group sizes and areas harvested. In essence, adaptive agents in our model solve the tradeoff between forest productivity and forest stress by reorganizing the structure of swidden labor networks (19). This form of adaptive self-organization leaves a power-law signature in the swidden mosaics because human decisions about field placement and field size relate to underlying spatial dynamics of natural forest seed dispersal, disturbance and regrowth processes. This occurs because field synchronization creates preferential-attachment dynamics (46), where existing swidden fields become the hub for placement of subsequent new fields. Fields require time to regain fertility before they can produce crops, so new synchronized fields are non-overlapping and expand the area in relation to the size and number of the synchronized fields attached to the original or hub field. This creates classic “rich-get-richer” dynamics where a small number of patches have many connections to other synchronized fields, while a larger number of isolated fields have few or no synchronized patches. The inflated number of highly connected swidden patches creates the “fat tail” and overall functionally-varied distribution of patches that are characteristic of scale-free power law distributions (11). Importantly, the overall size of these synchronized swidden patches act as larger forest disturbances and their spatial properties – indexed by patch size and correlation length *ε* in our analysis – relate to seed dispersal dynamics.

But do these patterns result from human activity alone, or can they be generated solely from the ecological dynamics of the model? To investigate this question, we explore how our model relates to other models of self-organization and criticality in biology and physics (11, 47–52). To do this, we configure the model with social dynamics “turned off”: all agents attempt to clear a field of the same size *X* and location strategy *S*. This eliminates all interactions among agents—there is no helping behavior, selective imitation, or decision-making. Consequently, *X* and *S* are treated as exogenous parameters. Despite this simplification, the results indicate that power-law distributions are observed across a wide range of parameters *X* and *S* (see “A simplified mechanistic model of swidden agriculture” section of the SM).

In light of this, what necessitates the use of a more complex model that accounts for the social dynamics of swidden groups? Both models generate patchy landscapes and power-law distributions of patch sizes, however the simplified model does not realistically capture the adaptive capacity of swidden agriculture that results from feedback between environmental conditions and the organization of labor and field placement strategies. In contrast, the simplified model assumes uniform agent behavior and ignores the adaptive nature of real-world decision-making based on trade-offs between costs and benefits, beliefs, social norms, and social interactions. Furthermore, it omits key social dilemmas faced by swidden communities, particularly the common-pool resource dilemma and the reciprocity dilemma, where cooperation in labor sharing can paradoxically lead to resource over-exploitation. However, the simplified model is useful for understanding how ecological disturbances can generate emergent landscape patterns. The environmental model most closely resembling our simplified model is one describing the dynamics of Kalahari vegetation (52) because both models incorporate local and global effects on landscape transitions between vegetated and cleared states. In the Kalahari model, non-linear transitions between vegetated and cleared states are driven by interactions between local ecological facilitation from seed dispersal and a global constraint that relates the proportion of tree cover with mean annual rainfall. In our simplified model, local effects also arise from ecological facilitation that is driven by seed dispersal, while global effects are driven by anthropogenic disturbance.

Our model also relates to the even simpler Ising model of ferromagnetism, which has been widely used to study phase transition phenomena in physical and social systems, including the Bali study that inspired our research (11). Several characteristics of the Ising model are analogous to both the simplified and full swidden models. Cells in both models can exist in only one of two states; the probability that a cleared cell becomes forested reflects the tendency towards magnetic alignment; field placement, synchronization, and helping behavior relate to temperature; and ecological facilitation acts as an external field. Critically, the Ising model can display completely different phase transition behaviors even under small changes, such as the introduction of random external fields or long-range interactions, and the resulting patch size distributions exhibit power laws (53). For a discussion of the relationship between the Ising model and the models presented here, see the section “Relationship to the Ising model” in the SM.

Overall, our main model explains the emergence and coevolution of two interlinked forms of self-organization. The first is social self-organization, driven by human behavior and interactions that resolve social dilemmas and promote sustainable levels of forest disturbance. The second is ecological self-organization, driven by anthropogenic disturbances structured by social self-organization, and resulting in power-law land-use patterns. Overall, our results support the hypothesis that the presence and scale of adaptive self-organization can be detected in the correlation of patches in couple human and natural systems. Together, our modeling and validation offers a unified picture of swidden agriculture as an adaptive self-organizing socioecological system, disentangling the intertwined social and ecological processes that shape swidden land-use dynamics.

## Conclusion

Until now, most models of swidden agriculture have remained mute with respect to how small-scale communities and villages could adapt to large scale fluctuations in complex systems like forests, which are subject to rapid and unpredictable change. Our model explains how the social norms and ecological knowledge of swidden farmers in small communities, like those in southern Belize, can reconcile emergent dynamics of social self organization and ecological self organization of forests to generate sustainable intensive swidden cultivation that enhances both harvests and ecosystem productivity at much larger spatial scales. These dynamics occur because swidden farmers frequently adapt their farming strategies to changing social and environmental conditions, which creates higher-level emergent effects on landscape dynamics involving seed dispersal and forest regrowth. The model can generate a range of realistic outcomes including sustainable cultivation with a range of spatial patterns, as well as deforestation due to overuse, and non-linear dynamics which create bifurcations points in ecosystem responses that cause forests to collapse due to human behavior or environmental stochasticity. Given the realistic outcomes produced by our model, along with statistical evidence from 18 societies worldwide, the model is likely generalizable to many societies where swidden agriculture occurs.

If correlated patchy environmental are indication of adaptive self-organization not only in Bali, but in highly varied swidden societies worldwide, stabilization of anthropogenic environments at a scale-free distributions of correlated patches may be a frequent characteristic of human-environment interactions in coupled human and natural systems. This raises important questions about the nature and extent of coupling that is required for adaptive synchronization. For example, does it occurs in other land use systems, including hierarchically organized societies? What is the range of temporal and spatial scales at which adaptive coupling can occur? And, can coupling occur in more complex land use contexts (e.g., contemporary urban–suburban-rural interfaces; pastoral communities in semi-arid zones where patchy mosaics of degraded, recovering, and intact lands; or managed wetlands and floodplains with overlapping zones of rice cultivation, fishing, and forests?) Our model represents the evolving coupled dynamics of small communities and their local environments, and our results indicate that sustainable swidden cultivation can occur under a wide range of environmental, social and demographic parameters. In instances where external drivers including market forces, land tenure regimes, or climate change undermine adaptive capacity and decease the resilience of swidden societies, our model demonstrates the remarkable flexibility of social norms for enabling communities to adapt to a wide range of internal and external pressures. Indeed, this flexibility helps explains the ubiquity and persistence of swidden in the Holocene. Our model also reveals a wider range of options for planners engaging with customary social norms and practices related to swidden that can support local management of community lands, thereby helping to avoid one-size-fits-all policy solutions, for example when engaging Indigenous and small-scale societies in the global fight against climate change (54).

## Materials and Methods

### Remote sensing data and classification

The study analyzes 18 swidden sites in Central and South America, equatorial Africa, central and southeast Asia, and Indonesia (Figure 1a). Each site was identified as using swidden practices from ethnographic literature and other publicly available sources (see Table S5 in the SM). We used Planet Lab’s Planet Explorer to search for documented place names and then identified swidden use areas visually to establish 50*km*^2^ circular areas of interest centered on the closest settlement where we downloaded standardized 4-band (RGB+NIR) remote sensing imagery with 3.125*m* (9.77*m*^2^) pixel resolution (55). Classification proceeded in six steps: First, we manually digitized training data for each site with QGIS (56), and using 13 classes (bare ground, recent clearing, early fallow, late fallow, mature forest, village/road, water, clouds, miscellaneous mask, palm oil fields, cattle pasture, urban, and wetland). Second, we used a random forest classifier (57) and segment anything (58) to classify the raster images. Third, we created a swidden raster with a subset of pixels that included the five classes directly related to swidden activity (bare ground, recent clearing, early fallow, late fallow, and mature forest), and set all other pixels to NULL. Fourth, we defined a synchronized swidden patch as a connected region containing two or more patches with the same functionally significant landuse class. In the 3.125*m* swidden raster imagery, individual swidden patches can be identified visually. To determine synchronized swidden patches, we down-sampled the swidden rasters using the raster::aggregate function, which applies a moving window and modal aggregation to create a coarser 46.875*m* pixel resolution (2, 197.27*m*^2^). This function effectively merges two or more adjacent patches that have the same land-use class into a single larger patch that is assigned the modal class, under the assumption that narrow buffers of different land types between contiguous patches have less functional significance with respect to successional dynamics and landscape ecology (Figure S9 in the SM). Fifth, to be able to compare the synchronized swidden raster with the simulated rasters from the model, the five swidden-related land use classes were reduced to two classes: forested (1, consisting of late fallow and mature forest) and empty (0, consisting of bare ground, recent clearing, and early fallow) pixels. Sixth, the area of each the merged patch in the synchronized swidden raster is then calculated by multiplying the number of pixels it contains by 2, 197.27*m*^2^ using landscapemetrics package (59). The resulting classified swidden mosaics are shown in Figures S4-S5 in the SM and the raster dataset used in the analysis is included in SM. The synchronized swidden rasters and the quantitative measurements of synchronized swidden patch areas are used in the statistical analysis.

### Statistical modeling

The statistical approach used in this study focused on testing for the existence of power laws in synchronized swidden patch size distributions because self-organization has been proposed as a plausible theoretical model for explaining spatial clustering in natural (52) and couple human and natural systems (11). Kolmogorov-Smirnov goodness-of-fit tests are used to assess whether a discrete power law is a plausible model for the patch size data extracted from the synchronization raster (60). We used a fixed *a*_*min*_ = 5 cells (corresponding to an area of 1.1 ha) and estimated the discrete power law scaling parameter *α* using maximum likelihood estimation as described in (60), along with the error estimate *θ* using a bootstrap procedure. We also tested two alternative models - discrete log normal and exponential using the Vuong likelihood ratio test for model selection (60, 61) (see Figures S6-S8 in the SM). In all cases, there is no evidence that either alternative provides a better fit than the power law model, so power law models alone were used to analyze simulation results. Analyses were performed in Matlab (Version R2024a) and in R employing the PoweRlaw package (62).

### Correlation function and correlation length

We adapt methods described in (11, eq. 1 and eq. 2) to analyze the spatial dependencies among swidden land-use classes by defining a correlation function that is based on the mutual information shared between all pairs of pixels in the synchronized swidden mosaic. Correlation function *C*(*d*) captures how much knowledge of the swidden class of one pixel reduces the uncertainty of knowledge of the swidden class of another pixel at a distance *d* from each other:

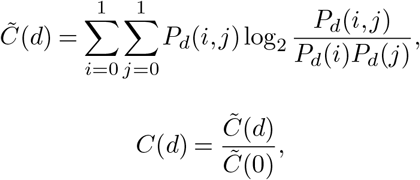

where *i* and *j* denote swidden land-use classes 0-1. *P*_*d*_(*i, j*) is the probability of observing vegetation classes *i* and *j* at the distance *d* from each other, *P*_*d*_(*i*) is the probability of observing a pixel of class *i* at the distance *d* from any other pixel. Note that if the vegetation mosaic is random, classes of pixels at a distance *d* apart are independent, resulting in a correlation function close to zero. Higher values of the correlation function correspond to stronger synchronization of swidden vegetation classes at a particular distance *d*. Overall, the correlation function *C*(*d*) is proportional to the Kullback–Leibler divergence between the actual distribution of swidden vegetation classes at distance *d* from each other and their distribution as if these classes were independent. We use the Manhattan distance to measure the distance between pixels. Given that each pixel in the synchronization raster is 46.875 m by 46.875 m, a Manhattan distance of 1 then translates to the distance of 46.875 meters. Based on the correlation function, we compute the correlation length as:

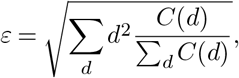

which captures the average synchronization distance in meters. Additional results on spatial patterning related to swidden field synchronization for all 18 sites are presented in Figures S4-S5 in the SM.

### Spatial agent-based modeling of swidden socioecological system

Additional details on the models used in this study are provided in the “Additional details on the model” and “A simplified mechanistic model of swidden agriculture” sections of the SM. The simulations were performed in Matlab (Version R2024a).

## Ethics Statement

Permission to conduct the ethnographic research that informed the model was obtained from village leaders according to customary cultural practices. Ethnographic research permits were obtained from the Belize Institute for Social and Cultural Research (ISCR/H/2/211). The research project was reviewed and approved by The Ohio State University Institutional Review Board (Study Id: 2017B0387).

## ACKNOWLEDGEMENTS

Many thanks to Steve Lansing and Sergey Gavrilets for providing comments on the manuscript. We are grateful for research funding provided to S.S.D. by the National Science Foundation USA (#1818597) and by The Ohio State University. Sergey Gavrilets at the University of Tennessee, and the College of Arts and Sciences at The Ohio State University provided high-performance computing support.

## Footnotes

1 Alpha values around 1 indicates a mixture of a small number of large forest patches, with a larger number of medium and small patches. Higher alpha values approaching 2 would indicate fewer large patches and more smaller patches

## Notes

### Competing Interest Statement

The authors have declared no competing interest.

